# Information and genetic counselling for psychiatric risks in children with rare disorders

**DOI:** 10.1101/542423

**Authors:** Andrew Cuthbert, Aimee Challenger, Jeremy Hall, Marianne van den Bree

## Abstract

**Purpose:** Genomic medicine has transformed the diagnosis of rare developmental disorders. Evidence about risks for co-occurring psychiatric disorders is, however, limited and potentially not fully considered when counselling families about a genetic diagnosis. Our study explored parents’ experiences of genetics services and their awareness of information concerning psychiatric risks.

**Methods:** Parents of children referred to paediatric services completed an online survey exploring, (i) experiences of receiving a genetic diagnosis; and (ii) sources of information about psychiatric, developmental and physical manifestations.

**Results:** Two-hundred and eighty-six respondents completed the survey. Thirty percent were unsatisfied with the communication of tests results. Higher satisfaction was predicted by personal communication (Odds Ratio 2.91); delivery by genetic specialists (OR 2.97); clear explanations (OR 5.14); and receiving support (OR 2.99). In contrast to developmental and physical challenges, psychiatric information was mostly obtained from non-professional sources, particularly for UK respondents (p <0.001). Information obtained from support groups was more helpful than from geneticists (OR 21.0), paediatricians (OR 11.0) and internet sites (OR 15.5).

**Conclusion:** The paucity of professional psychiatric information may compromise awareness of mental health risks for families. Further expansion of genomics in other services should integrate high quality genetic counselling training and provision of comprehensive mental and general health information.

## Introduction

Intellectual and developmental disabilities affect up to 1 in 25 children in high income countries.^1^ Clinical evaluation includes molecular cytogenetic investigations to improve diagnostic accuracy, inform genetic counselling, care management and ongoing family support. Barriers to identifying causal genomic variants in severely disabled children have substantially diminished as technological advances have translated into clinical services. Since their introduction in the mid 2000s, genome-wide tests have substantially increased the diagnosis of intellectual and developmental disorders, largely through detecting genomic micro-deletions and micro-duplications called Copy Number Variants (CNVs). First-line chromosomal microarray genotyping of children with intellectual and developmental disorders identifies pathogenic CNVs in approximately 10-15% of tests.^2^ More recent developments in clinical exome and whole genome sequencing have allowed further advances with the diagnosis of previously unexplained cases, bringing opportunities to improve diagnostic precision and offer new insights into disease mechanisms.^3–5^ These new developments however, require major changes in service delivery and clinical practice, including education and training for health professionals, service commissioning and patient engagement.^6^

CNVs in individuals diagnosed with developmental disorders are associated with increased risk of multiple psychiatric disorders, including Autism Spectrum Disorder (ASD), Attention Deficit Hyperactivity Disorder (ADHD), psychotic and bipolar disorders.^7–10^ For example, individuals with 22q11.2 Deletion Syndrome (22q11.2DS, formerly Velocardiofacial syndrome or DiGeorge syndrome) are at high risk of ASD, ADHD, anxiety, oppositional defiant disorder and schizophrenia.^11–13^ Emerging evidence concerning risks of neurodevelopmental, physical and psychiatric morbidity in diagnostic CNVs reported by clinical genetics services indicates a substantial variation in their penetrance and expressivity, which is susceptible to socioenvironmental modification.^14–16^ Despite these uncertainties, which currently limit personalisation of risk, treatment, prognosis and information provision, genomic testing is widely recognised as a major advance in health and social care for developmental disorders.^17^ Rapid increases in rates of genetic diagnosis and emergent knowledge concerning genotypic risk and phenotypic variability, impose significant challenges and considerations for genetic counsellors with responsibilities towards presenting accurate, comprehensible information while supporting patients and their families.^18–20^

In this study we explored parents’ opinions and experiences concerning psychiatric, neurodevelopmental and physical manifestations associated with genomic variants diagnosed in children with developmental disorders. Specifically, we asked parents about (i) their satisfaction with receiving genetic test results through attending specialist paediatric services; and (ii) the availability and practicality of information obtained about clinical manifestations of genomic disorders subsequent to their child’s diagnosis.

## Methods

### Survey respondents and procedures

We designed a 46 item online survey, using Online Surveys (https://www.onlinesurveys.ac.uk, Jisc, Bristol UK), for parents of children aged 0-17 years with developmental, intellectual and congenital disorders having a clinical genetic diagnosis. The survey comprised 4 main sections: (1) family demographics, child genotype, reported clinical diagnoses, family genetic history; (2) awareness of clinical disorders associated with the genetic diagnoses and sources of information used to understand them; (3) ratings of the quantity, content and helpfulness of information from each source; and (4) experiences of clinical services and receiving genetic test results.

An introductory section included information about the research team, the purpose of the study, instructions for participating, participant confidentiality and data protection. Participants provided consent by agreeing with the statement, “I have read the information above and on the previous page, understand that my participation is voluntary and I am happy to complete the questionnaire”. The study received ethical approval from Cardiff University School of Medicine Ethics Committee on 19 September 2014 (reference SMREC 14/34). Invitations to take part were distributed by advertisements in newsletters, websites and Facebook pages and by word-of-mouth at family support days sponsored by rare disorders support groups, including Unique – Understanding Rare Chromosome and Gene Disorders, Max Appeal, and Microdeletion 16p11.2 Support and Information UK.

### Statistical analyses

Response data was coded and downloaded from the survey website into the SPSS package (IBM Corp. SPSS Statistics for Mac, version 25.0, Armonk, NY, 2017) for statistical analysis. To test association between categorical variables in survey questions covering information sources and experiences of attending genetic testing services, 2 × 2 contingency tables were used and Pearson Chi-square tests. McNemar’s test was used to assess the significance of differences between correlated proportions in subjects assessed for pairs of binary variables, specifically to compare responses to questions about lay or professional sources of clinical information and questions about information utility. Odds ratios and 95% confidence intervals were calculated with VassarStats at vassarstats.net/propcorr.html. Wilcoxon’s signed-rank test was used to assess ordinal data for significance of differences between distributions of responses for non-independent samples. This was applied to Likert scale responses to questions concerning the amount and the content of information from multiple sources. Effect sizes (r) were derived from Z scores. Binary logistic regression was employed to test for association between demographic, genetic counselling and communication factors and satisfaction with genetics services. Specifically, the binary outcome variable was defined by parents expressing satisfaction (or dissatisfaction) with how their child’s genetic test result was communicated. The initial regression model incorporated family and child specific covariate using method ‘Enter’. Sequential hierarchical models incorporated genetic counselling specific and communication modality specific covariates. Where appropriate, ordinal variables with low responses for some categories were collapsed. Odds ratios correspond to the exponentiated unstandardised coefficients (beta weights) for each variable.

## Results

### Child and family characteristics

Two-hundred and eighty-six survey responses were recorded between 12 December 2014 and 31 May 2017. Of these 199 (61.6%) were located in the UK, 87 (26.9%) in the USA. Table 1 shows details of the respondents and children they reported on. The most frequent reasons for referral to paediatric genetics services were developmental delay (34.3%), congenital malformations (12.2%) and dysmorphic features (7.3%). There was extensive co-morbidity among children in the study, the mean number of diagnoses per child was 5.7. This included early developmental disorders (mean 2.4 per child); congenital malformations (mean 2.2 per child) and neuropsychiatric disorders including ASD, ADHD and Obsessive Compulsive Disorder (OCD), (mean 0.8 per child). In terms of genetic diagnoses, 127 (44.4%) reported unique CNV and 116 (40.6%), recurrent CNV associated with high risk of co-occurring neurodevelopmental manifestations. 39 respondents (13.6%) indicated their child’s CNV was inherited (table 1).

**Table 1.** Characteristics of the study sample and children diagnosed with genetic disorders. ^*1*^ Developmental Delay, Learning Disability, Speech and Language Delay. ^*2*^ Palatal, Cardiac, Respiratory, Musculoskeletal, Growth, Seizures/epilepsy, Sight, Hearing, Skin. ^*3*^ Attention deficit hyperactivity disorder, ADHD (N= 47), autism spectrum disorder, ASD (N= 86), obsessive compulsive disorder, OCD (N = 48), developmental coordination disorder, DCD (N = 49), dyslexia (N = 19). ^*4*^ Anxiety, Depression, and Schizophrenia or psychosis. ^*5*^ Recurrent neurodevelopmental susceptibility CNV loci: 1q21.1 deletion, 1q21.1 duplication, 2p16.3 deletion (NRXN1), 9q34.3 deletion, 15q11.2 deletion, 15q11.2 duplication, 15q11.2 (NOS), 15q13.3 deletion, 15q13.3 duplication, 15q13.3 (NOS), 16p11.2 deletion, 16p11.2 duplication, 16p11.2 (NOS), 16p12.2 deletion, 16p13.11 deletion, 16p13.11 duplication, 16p13.11 (NOS), 17q12 deletion, 22q11.2 deletion, 22q11.2 distal deletion, 22q11.2 duplication.

| Respondents |  |  | Children |  |  | Parent reported diagnoses |  |  |
| --- | --- | --- | --- | --- | --- | --- | --- | --- |
| Location | N | % | Age range (years) | Mean | Range | Clinical | N | Mean |
| UK | 199 | 69.9 | Age when diagnosed | 3.7 | 0-17 | Developmental<sup>1</sup> | 695 | 2.4 |
| USA | 87 | 30.4 | At time of survey | 8.5 | 0-42 | Physical or neurological<sup>2</sup> | 638 | 2.2 |
| Gender | N | % | Gender | N | % | Neuropsychiatric<sup>3</sup> | 230 | 0.8 |
| Female | 270 | 94.4 | Female | 125 | 43.7 | Mood and Psychotic<sup>4</sup> | 80 | 0.3 |
| Male | 16 | 5.6 | Male | 161 | 56.3 | Total | 1643 | 5.7 |
| Relationship | N | % | Reason for referral | N | % | Genetic data | N | % |
| Biological parent | 267 | 93.3 | Developmental delay | 98 | 34.3 | Neurodevelopmental CNV<sup>5</sup> | 116 | 40.6 |
| Adoptive parent | 10 | 3.5 | Congenital anomaly | 35 | 12.2 | Other CNV | 127 | 44.4 |
| Legal guardian | 2 | 0.7 | Dysmorphic features | 21 | 7.3 | Multiple CNVs | 16 | 5.6 |
| Other | 7 | 2.4 | Growth or stature | 16 | 5.6 | Named gene | 17 | 5.9 |
| Occupation | N | % | Neurological | 16 | 5.6 | Named syndrome | 5 | 1.7 |
| Employed | 141 | 49.3 | Multiple concerns | 13 | 4.5 | Unspecified | 5 | 1.7 |
| Carer | 78 | 27.3 | Learning disability | 11 | 3.8 | Familial CNV (parental) | 39 | 13.6 |
| Home maker | 58 | 20.3 | Family history | 10 | 3.5 | Siblings with CNV | 20 | 7.0 |
| Other | 9 | 3.1 | Neurodevelopmental | 10 | 3.5 | Other relatives with CNV | 13 | 4.5 |
|  |  |  | Neonatal concerns | 9 | 3.1 |  |  |  |
|  |  |  | Foetal anomaly | 7 | 2.4 |  |  |  |
|  |  |  | Metabolic | 2 | 0.7 |  |  |  |
|  |  |  | Unspecified | 38 | 13.3 |  |  |  |

### Experiences of genetics services

The majority of respondents received their child’s genetic test result from their paediatrician (130/286, 45.5%), geneticist (116/286, 40.6%) or genetic counsellor (28/286, 9.8%). Compared to the USA, parents in the UK were more likely to receive results from their paediatrician than from a genetic specialist (107/199 vs 23/87; p <0.001). Eighty-five respondents (29.7%) reported they were not satisfied with how their clinical specialist communicated the test result. Parents in the UK were more likely to be dissatisfied than in the USA (66/199 vs 19/87; *p* = 0.054). In terms of clinical specialism, parent satisfaction was more likely if results were communicated by genetic specialists compared to paediatricians (116/144 vs 78/130, p <0.001). This was significant for UK but not US parents (figure 1). Ninety-eight of 274 (35.8%) parents who received results from geneticists and paediatricians were not satisfied with the explanation given. Dissatisfaction was more frequent among UK parents (75/188 *vs* 23/86, p = 0.038), for whom geneticists were more likely to elicit satisfaction than paediatricians (p = 0.012, see figure 1). Ninety parents (31.5%) said they were not given information to accompany the genetic test result. More UK than US parents reported receiving test results without accompanying information (36.2% *vs* 20.7%, *p* = 0.009). Seventy-three percent (208/286) reported not receiving support when test results were delivered and 49.7% (142/286) recalled not being offered a follow-up appointment.

**Figure 1.**
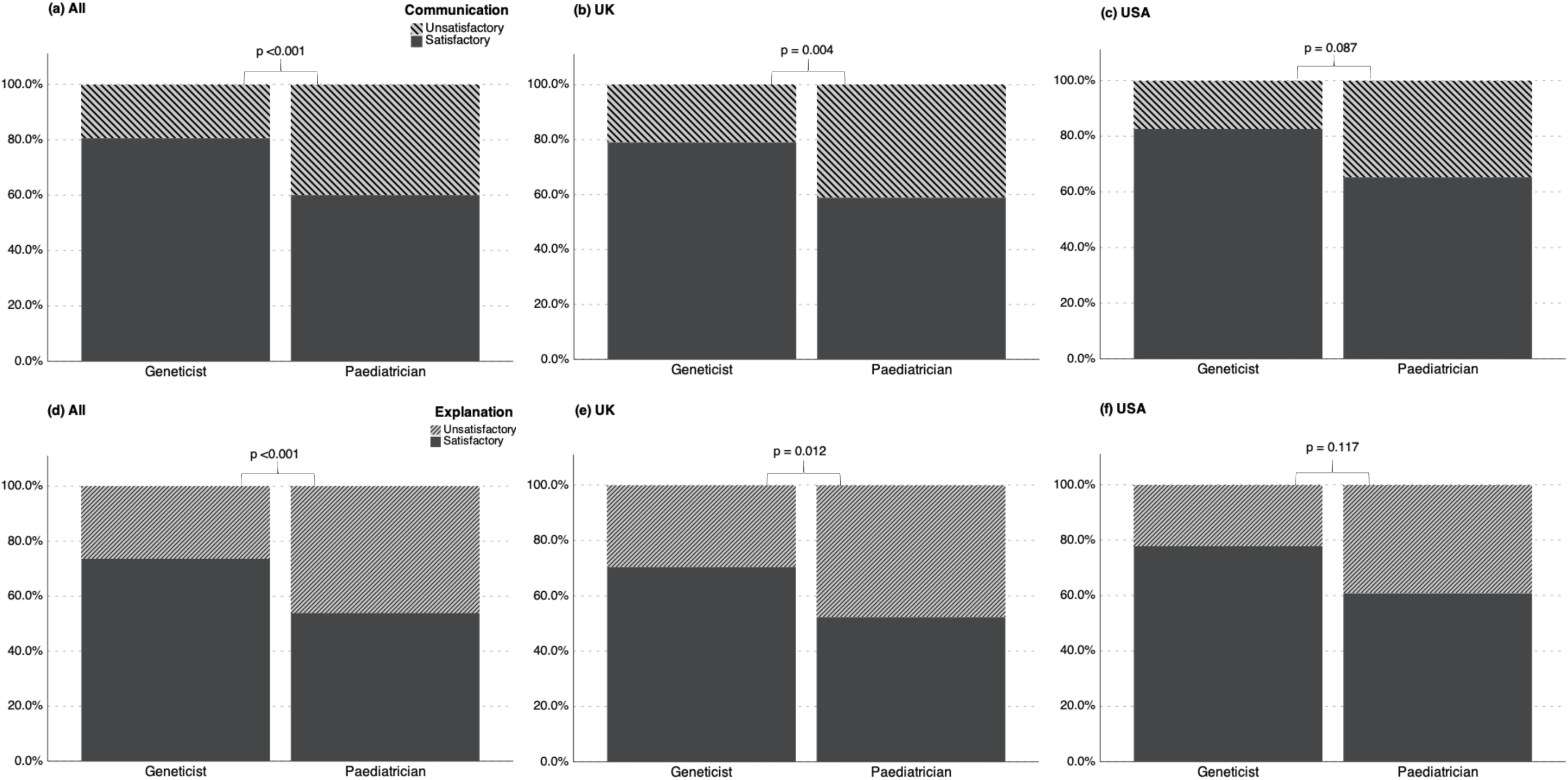
Comparisons of parent satisfaction with the communication and explanation of genetic test results by their child’s clinical specialist. Upper panel - comparison of satisfaction with communication of genetic test results between paediatric and genetic specialists: (a) all respondents; (b) UK respondents (N = 188); (c) US respondents (N = 86). Lower panel – comparison of satisfaction with test result explanations between genetic and paediatric specialists: (c) all respondents; (d) UK respondents (N = 188); (e) US respondents (N = 86).

To identify factors capable of predicting satisfaction with the communication of genetic test results, we carried out multi-level regression with three categories of variables; (i) child and family specific; (ii) genetic counselling; and (iii) modes of communication (see supplementary information: table 1). Satisfaction was predicted by variables in all 3 categories, the most significant being satisfaction with the explanation of results (OR = 5.14, 95% CI 2.58–10.26); receiving results from genetic specialists *versus* paediatricians (OR = 2.97, CI 1.41–6.26); receiving results in person *versus* communication by letter or telephone (OR = 2.91, CI 1.41– 6.26); and receiving support (OR = 2.99 CI 1.21–7.36). Receiving information to accompany the result did not independently predict satisfaction. Interestingly, child gender – in favour of males – also predicted satisfaction (OR = 2.56, CI 1.28–5.14) for which odds increased as additional variables were added (supplementary information: table 1). The final model accounted for 40.5% of the variance in the outcome variable.

### Sources of information on genomic disorders

We asked parents where they first obtained information about a variety of phenotypic manifestations they associated with their child’s genetic diagnosis. We combined responses for each source of information across 4 groups of disorders: developmental, physical, neuropsychiatric and mood and psychotic (table 2). sixty-five percent of the combined information on neuropsychiatric, mood and psychotic disorders came from non-professional (lay) sources compared to 41% for combined information on developmental and physical disorders, which was more often provided by health professionals (*p* <0.001, see table 2). For anxiety, depressive and psychotic disorders, parents predominantly gained information from lay sources (71.4%) in contrast to 39.3% for developmental delay, intellectual disability and speech and language difficulties (p >0.001).

**Table 2.** Parent reported sources of information about developmental, physical and mental health manifestations associated with their child’s genetic diagnosis. Totals for each group of manifestations comprise cumulative frequencies for individual conditions: (1) Developmental = global developmental delay (N=274), intellectual disability (N=267), speech and language delay (N=257); (2) Physical = cardiac defects (N=153), palatal defects (N=97), seizures/epilepsy (N=166); (3) Neuropsychiatric = autism spectrum disorder (N=203), attention deficit hyperactivity disorder (N=130), obsessive compulsive disorder (N=140), developmental co-ordination disorder (N=115); (4) Mood and psychotic = anxiety (N=154), depressive (N=85), schizophrenia or psychotic (N=73). Figures in parenthesis are counts for parent-reported associations between their child’s genotype and individual manifestations. ^*1*^ percentage values are for cumulative counts for individual manifestations in each group.

| Information source | Developmental and physical manifestations |  |  |  |  |  | Mental health manifestations |  |  |  |  |  |
| --- | --- | --- | --- | --- | --- | --- | --- | --- | --- | --- | --- | --- |
|  | Developmental |  | Physical |  | Combined |  | Neuropsychiatric |  | Mood + psychotic |  | Combined |  |
|  | N | %<sup>1</sup> | N | %<sup>1</sup> | N | %<sup>1</sup> | N | %<sup>1</sup> | N | %<sup>1</sup> | N | %<sup>1</sup> |
| **Health professional** |  |  |  |  |  |  |  |  |  |  |  |  |
| With diagnosis | 318 | 39.8 | 149 | 35.2 | 467 | 38.2 | 121 | 20.6 | 45 | 14.2 | 166 | 18.3 |
| At follow-up | 70 | 8.8 | 38 | 9.0 | 108 | 8.8 | 38 | 6.5 | 12 | 3.8 | 50 | 5.5 |
| Other | 96 | 12.0 | 49 | 11.6 | 145 | 11.9 | 64 | 10.9 | 34 | 10.7 | 98 | 10.8 |
| Total | 484 | 60.7 | 236 | 55.8 | 720 | 59.0 | 223 | 37.9 | 91 | 28.6 | 314 | 34.7 |
| **Non-professional** |  |  |  |  |  |  |  |  |  |  |  |  |
| Internet sites | 167 | 20.9 | 103 | 24.3 | 270 | 22.1 | 163 | 27.7 | 106 | 33.3 | 269 | 29.7 |
| Support groups | 74 | 9.3 | 40 | 9.5 | 114 | 9.3 | 100 | 17.0 | 66 | 20.8 | 166 | 18.3 |
| Other sources | 73 | 9.1 | 44 | 10.4 | 117 | 9.6 | 102 | 17.3 | 55 | 17.3 | 157 | 17.3 |
| Total | 314 | 39.3 | 187 | 44.2 | 501 | 41.0 | 365 | 62.1 | 227 | 71.4 | 592 | 65.3 |
| **Total (all)** | **798** |  | **423** |  | **1221** |  | **588** |  | **318** |  | **906** |  |

We examined differences in the origins of information between parents in the UK and USA (figure 2). Combined responses for all clinical manifestations, showed that parents in the UK were more likely to receive information from lay sources (primarily internet sites and support organisations) than health specialists (UK 57.6% *vs* US 38.3%, *p* = 0.005). For information about developmental and physical manifestations, UK parents were more likely to use non-professional sources (45.7% *vs* 31.5%, *p* = 0.042). The largest difference concerned information about psychiatric manifestations, for which 73.4% of UK responses involved using non-professional sources, compared to 47.9% in the USA (p <0.001). For anxiety, depressive and psychotic disorders, 80.8% of UK responses indicated the use of lay sources compared to 52.4% in the USA (p <0.001).

**Figure 2.**
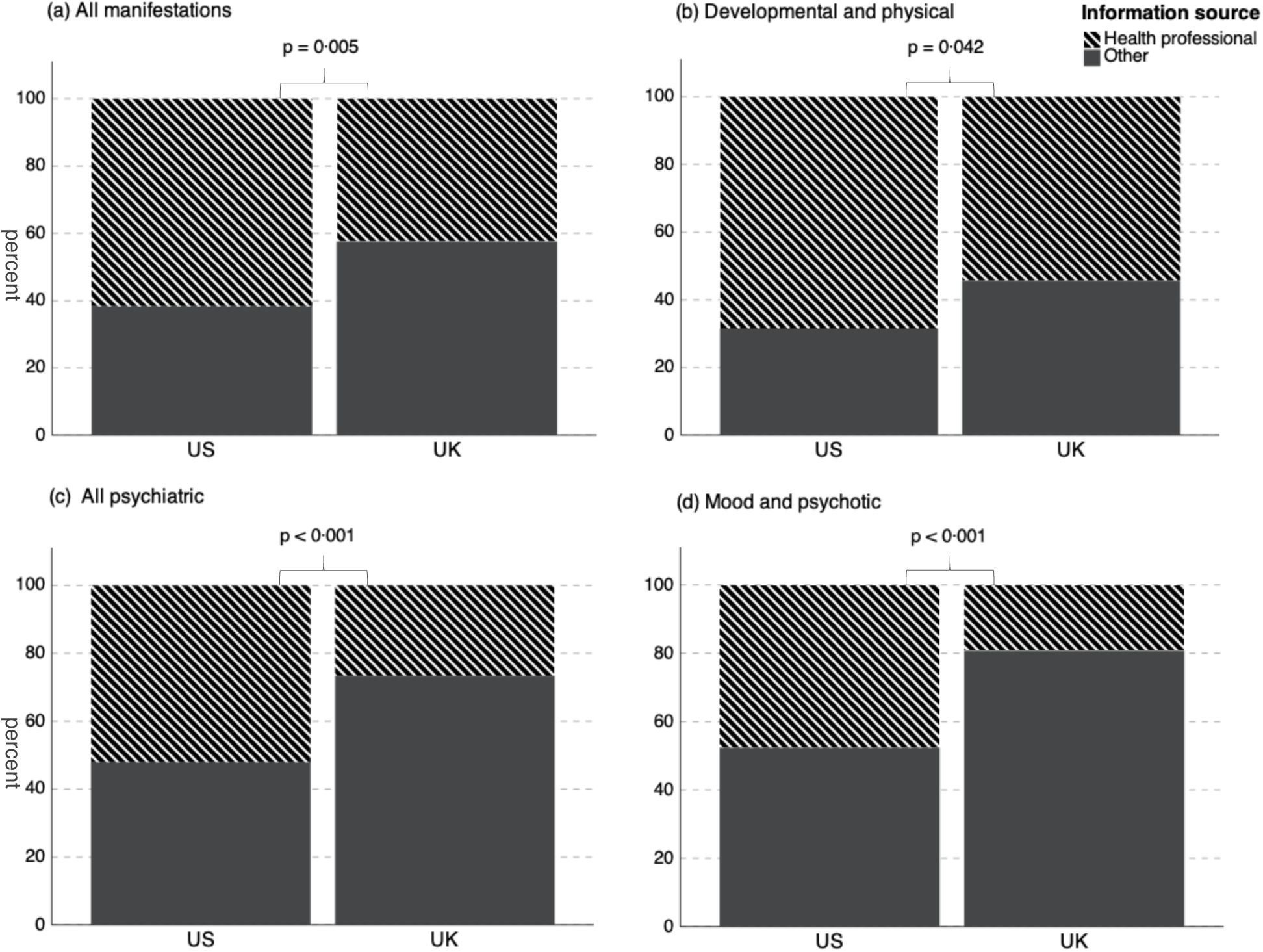
Comparisons of sources of information reported by respondents to learn about developmental, physical and psychiatric manifestations associated with rare disorders. Manifestations were grouped into developmental, physical, neuropsychiatric, mood and psychotic disorders (defined in table 2). Between country comparisons: (a) all manifestations; (b) developmental and physical; (c) all psychiatric (neuropsychiatric, mood and psychotic combined); and (d) mood and psychotic. Information sources: (i) Health professional sources = geneticist or paediatrician at time of diagnosis, at follow-up, or from another health professional; (ii) other sources = internet sites, support groups including Facebook, friends and family, books and leaflets.

We quantified the extent to which parents acquired information about recurrent manifestations of rare diseases from either professional, or non-professional sources, against typical developmental, physical and neuropsychiatric indications for paediatric referral; (i) intellectual disability (ID), (ii) cardiac defects, and (iii) ASD. In comparison to ID, parents were significantly more likely to gain information about all psychiatric manifestations from non-professional sources, with odds ratios ranging from 2.7 for ASD (95% CI 1.5 – 4.8) to 18.0 for depression (CI 2.4 – 134.8), whereas information about developmental delay was more likely to originate from health specialists (OR 13.0, CI 5.2 – 32.3, see supplementary information: figure 1). A very similar pattern was observed for all psychiatric traits when compared with cardiac defects (OR range 3.6 – 31.0). Finally, in comparison to ASD, information about anxiety (OR 16.0, CI 2.1 – 120.6), depression (OR 7.0, CI 1.6 – 30.8) and OCD (OR 6.0, CI, 1.8 – 20.4) was more likely to originate from lay sources, whereas information concerning physical and developmental conditions was more likely to originate from health specialists (supplementary information: figure 1).

We asked respondents to rate the amount, content and helpfulness of information available from 4 different sources (table 3). Amounts of information from support groups was rated as optimal (enough *vs* insufficient or excessive) more often compared to paediatricians (r −0.47, *p* <0.001), geneticists (r −0.51, *p* <0.001) and internet sites (r −0.38, *p* <0.001). Similarly, information from internet sites was more optimal compared to paediatricians (r −0.34, *p* <0.001) and geneticists (r −0.32, *p* <0.001). In terms of the content of information, support groups were again superior (clear and easy to understand *vs* too scientific or not relevant) to internet sites (r −0.31, *p* <0.001), geneticists (r −0.29, *p* <0.001) and paediatricians (r −0.14, *p* = 0.012). Likewise, parent-rated helpfulness of information from support groups was greater than internet sites (OR 15.5 [95% CI 3.71 – 64.77]; *p* <0.001), paediatricians (OR 11.0 [1.42 – 85.2]; *p* = 0.004) and geneticists (OR 21.0 [5.08 – 86.75]; *p* <0.001). Overall, information from lay sources was significantly more helpful compared to health professionals (OR 2.2 [1.37 – 3.53]; *p* = 0.001). Finally, between country comparisons revealed the content of information from support groups was more optimal for UK than for US parents (Z score −2.39, *p* = 0.02) as was the amount information from internet sites (Z score −1.95, *p* = 0.05).

**Table 3.** Comparisons of information obtained from health professionals and non-professional sources after receiving genetic test results. ^*1*^ Amount of information (too little; sufficient; too much) and ^*2*^ Content of information (too complicated; clear and comprehendible; not relevant) was assessed by pairwise comparisons of Likert scale responses; ^*3*^ The helpfulness of information (helpful *vs* not helpful) was assessed by testing differences in paired proportions; ^*4*^ Direction of effect, G = <u>G</u>enetic specialists; P = <u>P</u>aediatric specialists; I = Internet Sites; S = <u>S</u>upport groups.

| Comparison | Observations | Amount of information<sup>1</sup> |  |  |  | Content of information<sup>2</sup> |  |  |  | Helpfulness of information<sup>3</sup> |  |  |  |
| --- | --- | --- | --- | --- | --- | --- | --- | --- | --- | --- | --- | --- | --- |
|  |  | Z | r | p value | Direction<sup>4</sup> | Z | r | p value | Direction | Odds ratio | 95% CI | p value | Direction |
| Geneticists vs Paediatricians | 142 | -0.413 | 0.03 | 0.679 | G ≈ P | -1.196 | 0.07 | 0.232 | G ≈ P | 2.25 | 0.99 – 5.17 | 0.05 | G ≈ P |
| Internet sites vs Paediatricians | 182 | -4.522 | 0.34 | < 0.001 | I > P | -0.05 | 0.0 | 0.96 | I ≈ P | 1.88 | 0.80 – 4.42 | 0.144 | I ≈ P |
| Support groups vs Paediatricians | 120 | -5.17 | 0.47 | < 0.001 | S > P | -2.517 | 0.14 | 0.012 | S > P | 11.0 | 1.42 – 85.20 | 0.004 | S > P |
| Internet sites vs Geneticists | 360 | -6.037 | 0.32 | < 0.001 | I > G | -2.107 | 0.12 | 0.035 | I > G | 2.5 | 1.45 – 4.32 | 0.001 | I > G |
| Support groups vs Geneticists | 252 | -8.055 | 0.51 | < 0.001 | S > G | -5.336 | 0.29 | < 0.001 | S > G | 21.0 | 5.08 – 86.75 | < 0.001 | S > G |
| Support groups vs Internet sites | 330 | -6.957 | 0.38 | < 0.001 | S > I | -5.609 | 0.31 | < 0.001 | S > I | 15.5 | 3.71 – 64.77 | < 0.001 | S > I |

## Discussion

This is the first quantitative survey of its kind to investigate how parents experience the communication of genomic tests for rare developmental disorders and their subsequent endeavours to gain knowledge about the psychiatric risk implications associated with their child’s genetic diagnosis. This is an important topic; our survey indicates that parents had mixed experiences of attending genetics services. A large proportion of respondents in the UK reported dissatisfaction with genetic counselling for communication of results and indicated that the accompanying information they received from clinical specialists was less helpful than from other sources. In particular, parents relied extensively on non-professional sources for information about psychiatric risks. They often received sub-optimal information from clinical specialists, who tended to focus on early developmental and physical disabilities typically present at diagnosis. Families in the UK were more reliant on internet and Third Sector groups for clinically relevant information than in the USA, particularly information concerning psychiatric problems their children may be susceptible to.

The findings extend previous evidence showing that families often have difficulties obtaining satisfactory information from clinical specialists to aid comprehension of genomic tests results.^21–23^ We have revealed new evidence concerning a broad range of genomic disorders consistent with earlier findings concerning 22q11.2 deletion syndrome - one of the most frequently diagnosed genomic disorders - in which parents predominantly obtained psychiatric information from internet sites and support groups.^24, 25^ Consistent with studies of rare disorders more generally, we have shown that the great majority of parents search the internet to help comprehend their child’s diagnosis.^26^ However, many families do not seek medical information online, particularly in more deprived socioeconomic groups.^27^ Inadequate access to information is associated with increased stress and uncertainty for parents and negatively influences their engagement with services, potentially denying them the advantages of a precise diagnosis.^28^ Our survey did not gather sufficient demographic data to determine whether hard-pressed families were less likely to seek online content. However, we are concerned that socially disadvantaged families may not have the same degree of access to high quality information, support and services to which they are equally entitled.^29^

Paediatric and genetic specialists tended to focus on providing information about developmental and physical disabilities. This suggests that suitable information about psychiatric risks may be relatively scanty, or that addressing emotional and behavioural challenges is a lower priority when providing diagnostic genetic counselling. Alternatively, genetic counsellors may lack the requisite knowledge and skills to address psychiatric concerns. In turn, this may hinder information provision or referral to support agencies. Concerns expressed about overloading parents with information in the aftermath of a genetic diagnosis need to be balanced against the best interests of children and their parents.^30, 31^ Consistent with earlier studies of rare chromosomal disorders, our findings also reveal that parents often consider information from clinical specialists to have lower practical value than material found online or obtained from support groups and is frequently too complicated or irrelevant.^32^

Our findings reveal that patterns of information gathering differed between UK and US families. In the US, parents received a broader range of medical and mental health information from their clinician after diagnosis, whereas those in the UK were less likely to receive psychiatric risk information from health specialists. More than half of families in the UK had their child’s diagnosis communicated by paediatricians, compared to one-in-five in the USA. However, more seldom provision of professional psychiatric information in the UK was not explained by differences between clinical professionals who delivered results in each country. Our survey failed to uncover evidence explaining why psychiatric information is seldom offered by paediatric and genetics specialists in the UK, or conversely why those in the US offer a more comprehensive range of information. Conceivably, service-driven factors such as limited clinic time, or differences in professional development and training may account for observed differences. A clearer understanding of current limitations to providing comprehensive psychiatric genetic counselling when diagnosing developmental disorders would be welcomed and may reveal opportunities for resolving existing gaps in knowledge for concerned patients and families.

Opinions vary on the benefits and risks of online health information, ranging from concerns among health professionals over accuracy and regulation, to sociologists’ endorsements, praising its contribution to client empowerment. Between these views exists evidence that the general public adopt contingent behaviours towards online content, discriminating between trustworthy and untrustworthy content pages to supplement rather than replace traditional media.^33^ Importantly, near universal online search methods are increasingly concordant with the structure of internet health information and the hierarchical nature of results created by popular search engines. Therefore, we urge the development of new initiatives to explore how clinicians may improve access to information around the time of diagnosis, including signposting to high quality content beyond traditional media and assist Third Sector groups to innovate and procure new ways of supporting children with complex neurological disabilities.

The findings presented here are timely; paediatric genetics services strike a fine balance between delivering high quality services for escalating referrals and performing increasingly sophisticated tests with lengthy consent procedures. That a large proportion of families in our survey expressed dissatisfaction receiving genetic diagnoses, particularly in the UK, highlights the challenge facing long-established services. As the availability of genomic testing in mainstream clinical services increases, demands for relevant patient-oriented information concerning genetic influences on psychiatric risks are predicted to increase.^34, 35^ However, informing patients and families about complex and uncertain implications of genetic tests is challenging, requiring highly specialised skills in communication and psychosocial support. Personalising psychiatric risk across the lifespan of affected children such that service users understand the true nature of their risks, is likely to become a priority for psychiatrists and allied professionals working in the realm of rare disorders.^36, 37^ We recommend that additional education and training in brain disorders is prioritised for all clinical specialties offering genetic tests. Curriculum content for skills training should be developed accordingly, ensuring genetic counselling includes meaningful conversations about the full spectrum of medical and mental health risks and is accompanied by contemporary, relevant information.

Our study has limitations. Recruitment was biased in favour of parents who engage with the support groups who promoted the survey, potentially limiting the generalisability of our findings. The survey was designed with broad accessibility in mind. However, online surveys require significant internet literacy and time for participation, which may be difficult for some families. Parents who readily access the internet, are, conceivably, more likely to use it for seeking information related to the content of this study. The survey was overwhelmingly completed by mothers. However, families with severely disabled children are rare and not representative of the general population. This may be a reflection of the socio-domestic influences on caregiving in the context of disability.^38^ Also, we were unable to independently verify health information provided by parents. Self-report surveys have recognised shortcomings exacerbated by respondents potentially having to recall facts and experiences several years after the event. Despite these limitations, the views and experiences expressed by a large number of families provides a timely insight into the current state of medical health information, clinical services and lay support for the developmental disorders community.

In summary, our findings reveal that parents are inadequately informed by their clinical specialists about potential neurodevelopmental and psychiatric challenges in the context of paediatric genetic testing. Parents search extensively for information about their child’s genetic diagnosis, retrieving neuropsychiatric information primarily from internet sites and lay support groups, where its accuracy and validity is unregulated and potentially less reliable. We believe the current results are important in informing service development and training in both clinical genetics and psychiatry. In particular, they point to the need for closer integration of medical genetics and psychiatry to address the needs of those receiving a genetic diagnosis. Future initiatives should focus on identifying measures to promote the inclusion of psychiatric risk information, provide greater support in genetic counselling clinics and ensuring that education and training for practitioners to encompass the full spectrum of neurodevelopmental and mental health challenges faced by children with genomic disorders.

## Acknowledgements

We are very grateful to all the families who helped with this study. We thank Dr Beverley Searle and all the staff at the charity Unique - Understanding Rare Chromosome and Gene Disorders, for their support and for promoting the study. We also wish to thank Max Appeal for their assistance with the advertising the survey. We are grateful to Dr Michael Arribas Ayllon (School of Social Sciences, Cardiff University) for his valuable contribution. The work was supported by The Waterloo Foundation (Grant No. 506926) and the Medical Research Council (Grant No. MR/N022572/1).

