## Supplementary Information for "Information and genetic counselling for psychiatric risks in children with rare disorders"

**Supplementary table 1:** Multilevel logistic regression of predictors of satisfaction with receiving genetic test results. [1] Binomial regression with family specific variables only; [2] Hierarchical model with added genetic counselling specific variables; [3] Hierarchical model with added communication modality variables.

| | **Model A**[1] | | **Model B**[2] | | **Model C**[3] | |
|---|---|---|---|---|---|---|
| **Variables** | OR (95% CI) | *p* value | OR (95% CI) | *p* value | OR (95% CI) | *p* value |
| **Level 1 (child and family)** | | | | | | |
| Family Location (vs UK) | **2·30 (1·17–4·53)** | **0·016** | 1·62 (0·77–3·38) | 0·201 | 1·27 (0·57 – 2·87 | 0·564 |
| Child's gender (vs female) | **2·18 (1·21–3·93)** | **0·009** | **2·54 (1·31–4·91)** | **0·006** | **2·56 (1·28 – 5·14)** | **0·008** |
| Age at genetic diagnosis (years) | 0·93 (0·83–1·03) | 0·159 | 0·91 (0·80–1·02) | 0·114 | 0·90 (0·79 – 1·02) | 0·094 |
| Respondent's gender (vs female) | 0·0 | 0·998 | 0·0 | 0·998 | 0·0 | 0·998 |
| Home maker/carer (vs employed) | 0·60 (0·33–1·07) | 0·083 | 0·68 (0·36–1·31) | 0·251 | 0·57 (0·29 – 1·13) | 0·107 |
| **Level 2 (genetic counselling)** | | | | | | |
| Satisfactory explanation (vs not satisfied) | | | **5·47 (2·82–10·58)** | **<0·001** | **5·14 (2·58 – 10·26)** | **<0·001** |
| Offered information (vs not offered) | | | 1·22 (0·62–2·43) | 0·563 | 1·05 (0·51 – 2·16) | 0·896 |
| Offered support (vs not offered) | | | **2·83 (1·22–6·56)** | **0·015** | **2·99 (1·21 – 7·36)** | **0·017** |
| Offered follow-up (vs not offered) | | | 1·05 (0·55 – 2·01) | 0·889 | 1·15 (0·59 – 2·27) | 0·68 |
| **Level 3 (mode of communication)** | | | | | | |
| Given by geneticist (vs paediatrician) | | | | | **2·97 (1·41 – 6·26)** | **0·004** |
| Given in person (vs letter/telephone) | | | | | **2·91 (1·43 – 5·94)** | **0·003** |
| **Nagelkerke R-square** | 0·145 | | 0·335 | | 0·405 | |

**Supplementary table 2:** Pearson correlation matrix for variables used in the study of service experiences.

| Variables | Location | Parent gender | Parent occupation | Child gender | Age at diagnosis | Who gave result | How given | Satisfied with communication | Satisfied with explanation | Given information | Given support | Offered follow-up |
|---|---|---|---|---|---|---|---|---|---|---|---|---|
| Location | 1 | | | | | | | | | | | |
| Parent gender | 0.029 | 1 | | | | | | | | | | |
| Parent occupation | -0.003 | -0.105 | 1 | | | | | | | | | |
| Child gender | -0.092 | -0.03 | 0.003 | 1 | | | | | | | | |
| Age at genetic diagnosis | -0.002 | -0.004 | 0.009 | -0.037 | 1 | | | | | | | |
| Who gave result | **.301**** | -0.017 | 0.014 | -0.1 | 0.066 | 1 | | | | | | |
| How given | -0.064 | -0.066 | 0.049 | 0.119 | -0.111 | -0.039 | 1 | | | | | |
| Satisfied with communication | -0.114 | **.158**** | 0.037 | **-.167**** | -0.062 | **-.230**** | **-.224**** | 1 | | | | |
| Satisfied with explanation | **.123*** | -0.021 | -0.017 | 0.013 | 0.021 | **.203**** | 0.035 | **-.416**** | 1 | | | |
| Given information | **.153**** | -0.034 | -0.027 | 0.071 | -0.073 | 0.062 | **.135*** | **-.202**** | **.255**** | 1 | | |
| Given support | 0.039 | -0.09 | -0.034 | -0.03 | **-.128*** | -0.027 | **.210**** | **-.192**** | **.157**** | **.212**** | 1 | |
| Offered follow-up | 0.066 | -0.091 | **-.128*** | 0.024 | -0.08 | -0.054 | 0.041 | -0.048 | -0.034 | -0.02 | 0.101 | 1 |

** $p < 0.01$
* $p < 0.05$

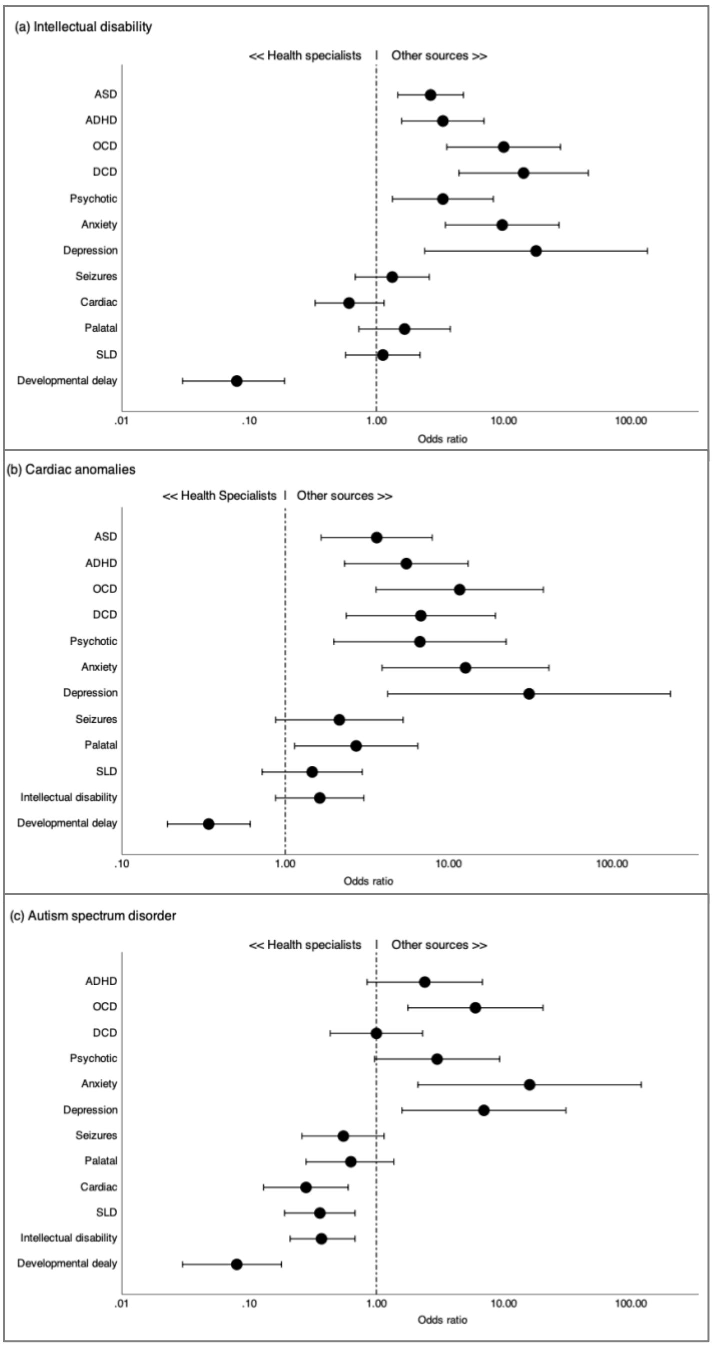


**Supplementary Figure 1:** Variation in information sources for individual manifestations. Dot plots depicts odds ratios for pairwise comparisons between sources first used for information about recurrent CNV-associated manifestations compared with sources representative of three major classes of disorder: (a) developmental disorders (intellectual disability); (b) congenital disorders (cardiac anomalies); (c) neuropsychiatric disorders (autism spectrum disorder). Sources: (i) health specialists = clinician at time of genetic diagnosis; clinician at follow-up appointment; other health professional, (ii) other sources = internet sites; voluntary support groups (including Facebook groups); friends and family; books and leaflets). Bars represent 95% confidence intervals.
